# Determining Interaction Directionality in Complex Biochemical Networks from Stationary Measurements

**DOI:** 10.1101/2024.04.16.589270

**Authors:** Nava Leibovich

## Abstract

Revealing interactions in complex systems from observed collective dynamics constitutes a fundamental inverse problem in science. Some methods may reveal undirected network topology, e.g., using node-node correlation. Yet, the direction of the interaction, thus a causal inference, remains to be determined - especially in steady-state observations. We introduce a method to infer the directionality within this network only from a “snapshot” of the abundances of the relevant molecules. We examine the validity of the approach for different properties of the system and the data recorded, such as the molecule’s level variability, the effect of sampling and measurement errors. Simulations suggest that the given approach successfully infer the reaction rates in various cases.

## I. INTRODUCTION

Many complex systems in physics and biology constitute networks of dynamically interacting units [1]. In these systems, especially in biological ones, the process is often described with stochastic variables [2–4], and their collective dynamics and functions are governed by the network structure. Determining the underlying interaction topology is essential, for example, for understanding and controlling their function, identifying new pathways in gene regulatory networks or understanding some metabolic mechanisms [5–8]. Particularly, understanding the direction of metabolic reactions is fundamental for predicting cellular behavior, optimizing metabolic engineering strategies, and identifying potential drug targets. Revealing these interaction networks poses a great challenge. Therefore, various studies have examined methods to find the structure of interaction networks [9–15].

Commonly, interaction networks are constructed from data obtained by time-series measurements, or even pseudo-time trajectories [12, 13, 16–23]. To do so, researchers use some mathematical and statistical tools such as Bayesian inference and maximum likelihood together with machine-learning algorithms [12, 13, 19–22]. There, the collective dynamics are used for network inference - whether the interactions are approximately linear or remain nonlinear, where the latter requires further assumptions on the system [6, 24, 25]. However, multidimensional time-dependent synchronized recorded data is not available in some scenarios. The use of steady-state data is also examined, yet the dynamics are assumed to be linear in the activity of the nodes of the network [26], or the system is assumed to be externally driven in a controlled way [25, 27]. Nevertheless, real systems are not required to fulfill these requirements.

In many cases the collective dynamics evolve via the (Markovian-) Master equation, where stochastic variables possess integer values [28]. Specifically, one of the well known models studied using the Master equation is the stochastic birth-death process which describes stochastic biochemical reactions [28]. In general, interaction networks of variables which evolve with Master dynamics are not limited to biochemical reaction networks, and has also been studied in other disciplines, see [29] and references therein. Yet, for consistency, this manuscript is oriented to bio-chemical reaction network studies, with adequate terminology.

In biochemical reaction network studies, one may infer the *functional* connectivity which gains insight into statistical dependencies that the entire set of interactions yield between pairs of units through the collective network dynamics [30, 31]. The statistical dependencies between components may be given by the well-known Pearson correlation, the mutual information, or their analyses with silencing methods [31–33]. These, however, are symmetric metrics and as such they are blind to the direction of the interaction providing only non-directed networks. Non-symmetric metrics, such as partial correlations and dependency analyses, require recorded data from all other interacting variables within the network [34–38]. That data is not available in many systems.

Here, we introduce a novel data analysis approach to infer the interaction direction, which under some conditions may be interpreted as causality. Its key advantage is that only the variables under investigation need to be measured, while the mechanistic details of regulation or degradation of all other components within the network do not need to be modeled or even to be observed, see Fig. 1. This approach exploits the global probability flux-balance equation that must be satisfied in the steady state [39–41]. In contrast to existing works, our approach does not require temporal information, experimental perturbations, or complete observation of all components within an interaction network.

**FIG. 1.**
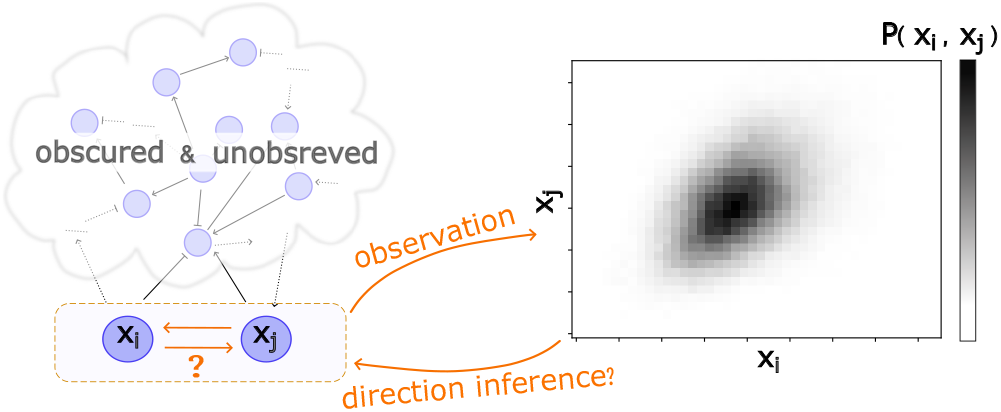
Main goal. Determining the direction of interaction between stochastic variables’ units is challenged by lacking of any temporal observations, any further knowledge on the directed network topology, or any records of the other variables within that network. We show that the steady-state joint distribution can be exploited to infer the arrow direction using the global balance equation Eq. (2).

## II. RESULTS

### A. Theoretical Background and Overview

Consider *V* variables coupled via an interaction network, i.e., a directed graph with *V* vertices. The variables state vector is given by 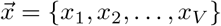. Each variable *x*_*i*_(*t*) ∈ N_0_ and evolves with time by probabilistic events 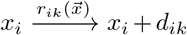 where the *k*-th reaction changes the abundance of the *i*-th molecule *x*_*i*_ by *d*_*ik*_. The reaction rates 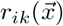 depend on the state vector 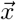 according to the interaction network. The stationary Master equation yields the global balance relation, which means that for every node *i*

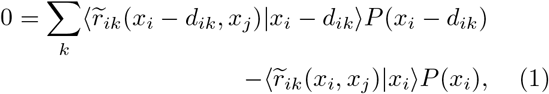

where 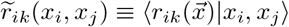 and the angular brackets represent conditional means. The above relation is derived from a summation of the Master equation over all other variables where consider stationarity *∂*_*t*_*P* = 0 [39–42], see further derivation in Methods and SI.

From Eq. (1) one may obtain the reaction rate only from a given joint steady-state distribution of the two molecules *P* (*x*_*i*_, *x*_*j*_), see Methods and [42]. Our approach is based on *local* sensitivity of 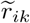 to *x*_*j*_; a direction of the interaction is inferred, which means an arrow is drawn in the directed graph, if a sufficient sensitivity is quantified, see details in Methods and SI. We remark that the suggested directional inference approach solely uses information about (*x*_*i*_, *x*_*j*_), thus the directionality inference problem decomposes over pairs of nodes within the network such that the direction of each interaction can be reconstructed independently.

To evaluate the quality of the inference, we initially consider a class of random networks with Michaelis–Menten kinetics which model gene regulatory networks [43], and random networks where each unit is a Goodwin oscillator, a prototypical biological oscillator that characterizes various biological processes such as circadian clocks [44, 45]. Additionally, we examine our inference strategy on a biological realistic system which is based on a subset of an *E. coli* gene regulatory network given in [46–48]. In the following, we evaluate the performance of the suggested approach to identifying the direction of a given interaction.

### B. Performance in General Synthetic Networks

Our strategy is successful in inferring the direction of the interaction between two nodes within an unknown network of interactions, where only the two-node joint steady-state distribution is given as the input see figures 2-4. It demonstrates that stationary probability distributions contain enough information to deduce the direction of a given interaction within an unknown network of interactions.

**FIG. 2.**
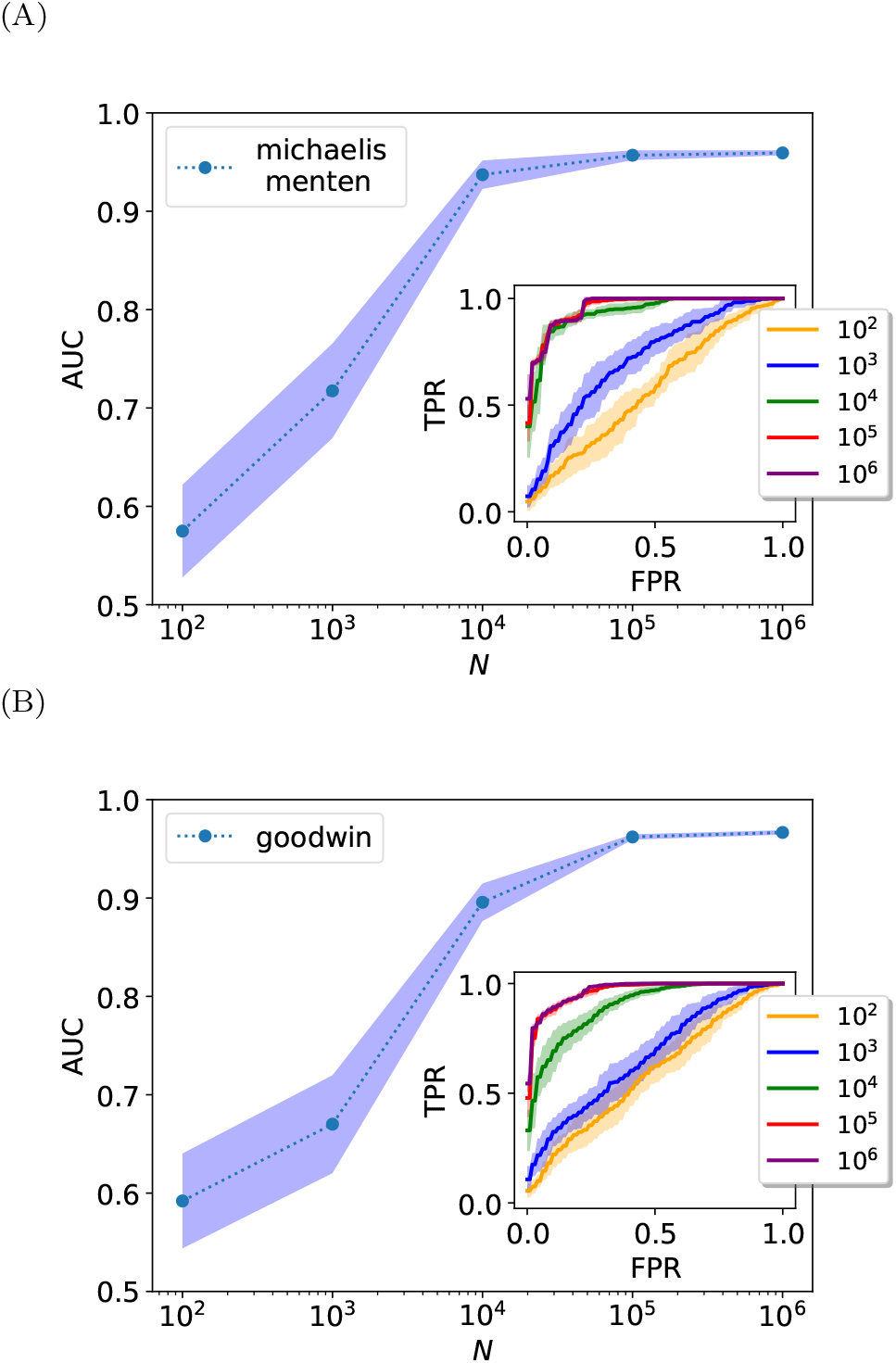
Inferring interaction directions from steady state. Simulation results indicate an improvement with increasing the number of data points *N*. Panel (A): The AUC scores for the Michaelis-Menten model (see methods). Inset: its corresponding ROC curves. Panel (B): Similar results are obtained for the coupled Goodwin oscillators. The shaded area corresponds to the standard deviation.

One of the most commonly used classifiers performance scores is the receiver operating characteristic curve (ROC) - which compares the true positive rate (TPR) to the false positive rate (FPR) at different threshold settings. A perfect classification yields TPR=1 with FPR=0, while a random one forms a diagonal line where FPR = TPR. The area under the receiver operating characteristic curve (AUC) is a single-valued score indicating overall performance, ranging from 0.5 for random to near 1 for perfect classification. We use these to evaluate the suggested direction inference strategy.

As aforementioned our method poses a good direction classifier - it successfully infers the direction of a given edge where the relevant joint distribution is sufficiently sampled. As is shown in Fig. 2 the direction inference improves with a larger set of data points. In particular, a sufficiently large data set provides scores beyond the random classifier. It signifies that the direction of interaction is indeed encoded in the stationary snapshot of many data points. Importantly, recall that the input is simply the two-variables joint distribution thus our method allows decomposition of the network - which means that within a given un-directed graph one can learn the direction of a single edge independently, regardless of other edges, see further discussion in SI. In the following, we discuss some key aspects that we found essential for the underlying goal. In particular, we examine some determinants that may affect probability to correctly infer directionality.

### C. Determinants and Limitations

The stationary joint probability density between *x*_*i*_ and all variables directly interacting with the *i*-th molecule must satisfy Eq. (1), hence it acts as a self-consistency test for stationarity. It is shown in [42] how this relation can be “inverted” to determine rate functions from observed probability densities. In other words, the information about the dynamical rate of a variable with its dependence on its coupled variables is indeed encoded within the joint steady-state distribution. However, while Eq. (1) must provably hold for all stationary states, empirically observed distributions carry sampling errors that can significantly impact inferred rates [42], leading to potential errors in determining the interaction direction. Our results reflect this errors dependence, as illustrated in Fig. 2; larger *N*, which is equivalent to lower sampling errors, yields increased accuracy (SI).

A further characteristic examined is the index of dispersion *D*, i.e. the variance to mean ratio. It quantifies the variable dependence on others, where *D ≈* 1 indicates that the variable’s dynamic is weakly dependent on other variables (SI). We show in Fig. 3 (A) that higher probability of success inference is found were *D* is far from 1. We note that a similar behavior was previously reported in [42]. For given edge *x*_*j*_ *→ x*_*i*_ the functional dependence of 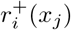 in the level *x*_*j*_ is better determined for higher mutual information between these two variables. Moreover, some local properties of the network may affect the performance of our direction inference method. We examine the in- and out-degrees, and the lifetimes of the molecules. Recall that we evaluate the sensitivity of 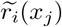 to *x*_*j*_, where 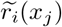 is a quantity averaged over all other income edges, thus we expect that the connectivity nature possesses an influence over the performance.

**FIG. 3.**
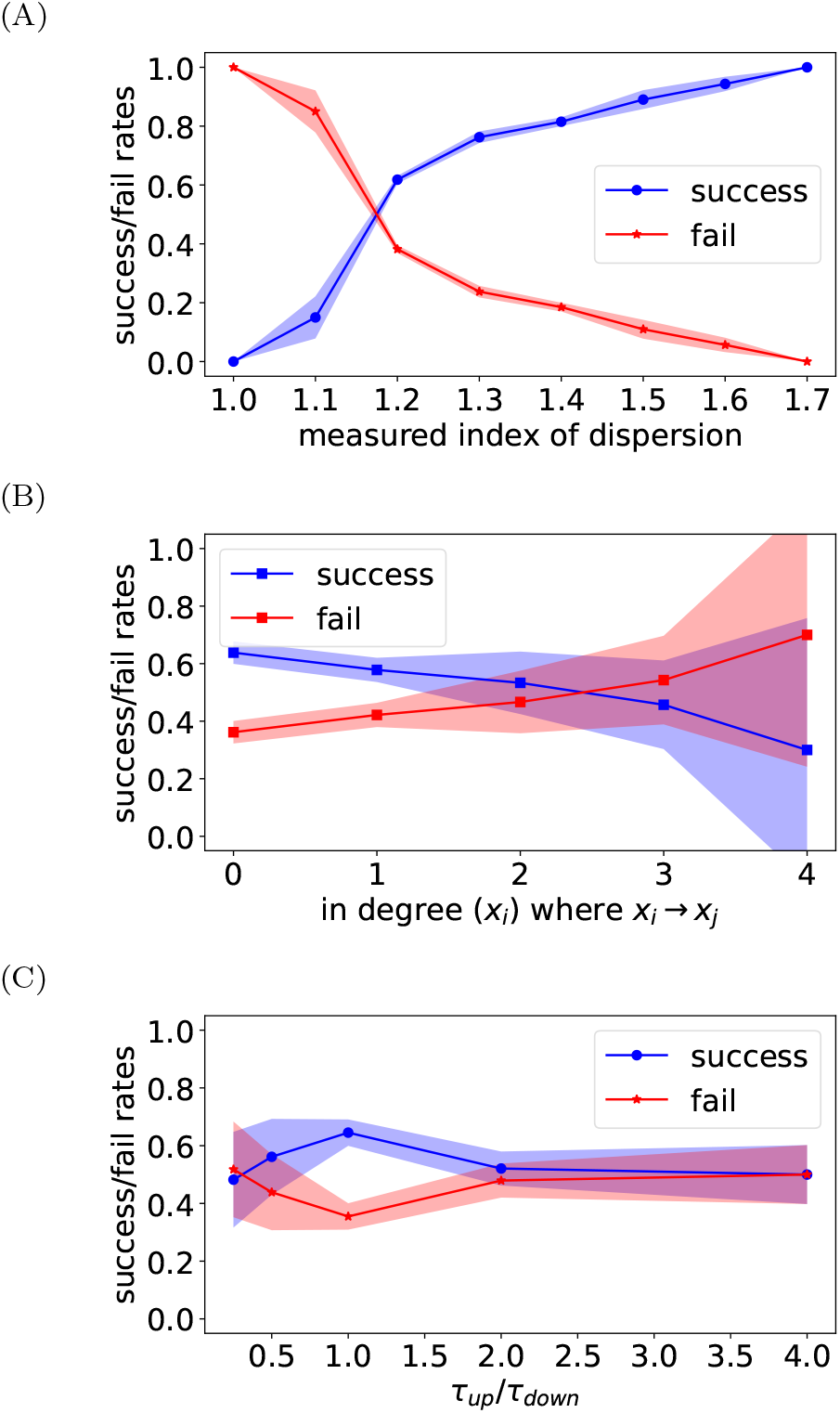
Factors dictate the performance. (A): the index of dispersion affects the probable for a successful inference of the direction. Results obtained the Michaelis-Menten model with *N* = 10^2^. (B) Dependence of the degree on the directionality inference and (C)) lifetimes dependence obtained from the Michaelis-Menten model with *N* = 10^2^. For all panels the shaded area represents the standard deviation

Additionally, the lifetime of a molecule is also expected to play an important role in the overall performance as is discussed in [42]. In the results presented in Fig. 3 (B) we have found that the in-degree of *x*_*j*_ presents some dependence over the performance, yet we did not find it significant, see further analysis and simulation results in SI. Moreover, we show in Fig. 3 (C) that a lifetimes ratio far from 1 results in decreasing the probability for an accurately inferred direction of influence, which agrees with [42]. Nevertheless, we comment that our network ensemble does not cover the entire possible network topologies space, an important factor that might skew the results, thus the nature of these phenomena needs further research.

### D. *E. coli* Gene Regulatory Network

We aggregate the insights from the above, and apply our strategy for the direction inference on a previously proposed biologically realistic system - a model which is based on *E. coli* gene regulatory network provided in the DREAM challenge [46–48], see Methods. The system shown in Fig. 4 poses various characteristic lifetimes and vertices’ degrees. We decompose the network and analyze each edge separately with its measured two-node joint distribution. As shown our inference method performs well for edges from nodes with zero in-degree with relatively long lifetimes (all edges from nodes 0 and 5), while erroneous direction emerges at edges connecting high-degree nodes with short lifetimes (nodes 1, 6 where each connected to two wrongly drawn arrows).

**FIG. 4.**
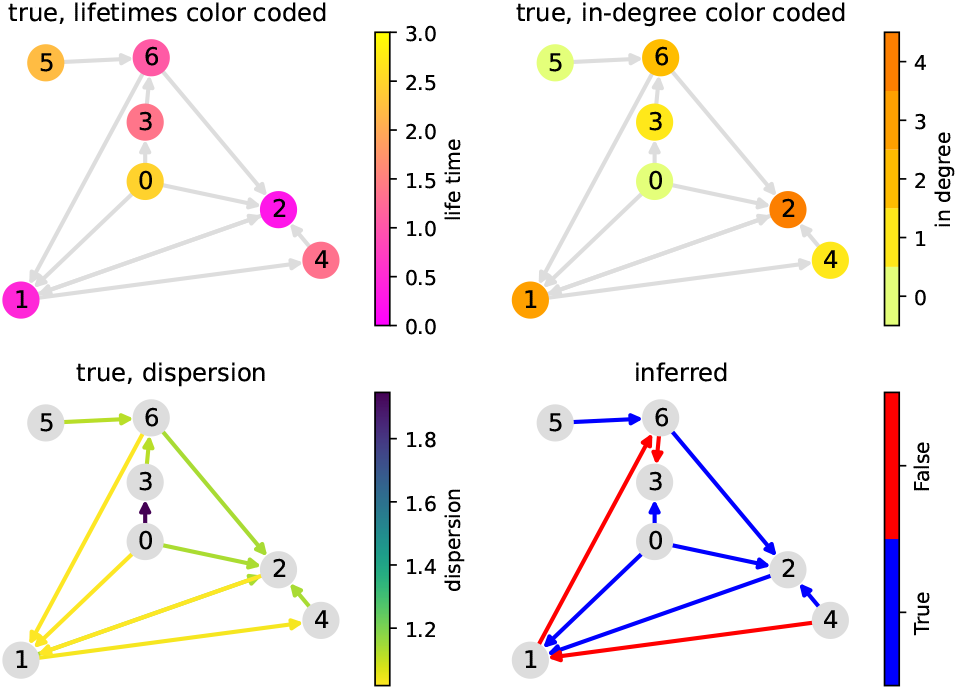
Simulation results for the *E. coli* gene regulatory network model.

## III. DISCUSSION

Unveiling the interaction network among stochastic variables and delineating the direction of each interaction is crucial for understanding biological systems. Here we present a method designed to infer the directionality within this network solely from the steady state of pertinent molecules. We assess the effectiveness of this approach across various properties of the system. Simulation results indicate that the proposed method demonstrates success in inferring the reaction direction in various cases.

As mentioned, our strategy relies on the time-invariant density, and as such possesses a benefit as explained in the following. Recorded data points can be employed without a known temporal order, taken at different sampling intervals, or aggregated from multiple experiments, as long as they were conducted under the same conditions. Hence this stationarity assumption enables the analysis of a wide range of biological systems. Importantly, Eq. (1) is not an approximation but corresponds to an exact flux balance relation at stationarity. The only dynamics excluded from our analysis is transient behavior such that stationary probability distributions are not accessible from experimental data. Even so, explicitly time-varying systems that technically never reach a stationary state, such as deterministic oscillations, may satisfy a similar form of Eq. (1) when considering their time-averaged probability distributions and rates. However, care has to be taken when comparing such time averages with population averages, see further discussion in SI.

The static snapshot provides statistical features of the stochastic variables, enabling the inference of the interaction direction. Statistical information is frequently used to gain insights from snapshots about the dynamics and the relations between variable components [27, 31, 32, 49–51]. Our strategy for revealing interaction directions thus constitutes a previously unknown intermediate approach, placed between purely statistical methods used to infer effective connectivity (such as correlations) and approaches that infer physical connectivity from high-resolution, time-ordered recordings of the complete dynamics [24–26, 52, 53]. In particular, statistical methods for inferring *directed* edges that rely on partial correlation analyses were applied for revealing directed graphs [35, 38, 54, 55]. These methods, however, require measurements from other components within the system and in that sense are non-local. Nevertheless, we tested the performance of the partial correlation analyses over our synthetic data and found that it is underachieved and thus inadequate for our goal (see SI).

Using the sensitivity analysis presents an equivalence with the examination of the slope of 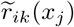 or its linear response. This linear sensitivity analysis for the interaction network identification is conceptualized and demonstrated for stationary systems under small perturbations [27]. There, the authors assume continuum variables that evolve via coupled differential equations with no noise or with additive noise. Conversely, our method considers discrete variables that propagate following the Master equation which effectively poses multiplicative noise. In addition, we note that our approach does not require externally driving the system in a controlled way that is not applicable in many systems [27].

Another possible directionality classification quantity is related to the Sum-of-Squared Errors (SSE), which might be associated with the Minimum Description Length (MDL). There the SSE (or the MDL) comparison between the two directions is well mathematically established [56–58]. Nevertheless, we show (SI) that our directionality classification quantity based on |*J*_*ij*_| out-performs the previously proposed SSE-based one, especially in low *N*. In the SI we further discuss the classifying quantities, with additional simulation results.

Harnessing of the global flux balance Eq. (1) to gain insight on the interaction quantification has been suggested recently [42]. There the authors demonstrated that the steady state joint distribution indeed encapsulates information about the dynamical features such as the birth rates. They assume that the direction of the edge, together with the stationary *P* (*x*_*i*_, *x*_*j*_), are given, and thus find the functional shape of 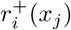 for various values of *x*_*j*_. Here we exploit this approach, but instead of looking for the full functional shape we only examine a partial behavior of 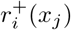 - its sensitivity analysis - without assuming the direction of the edge at all, see further discussion on SI.

Furthermore, we have examined some of the performance determinants, i.e. the properties which affect the efficacy of the direction’s identifier. For example, we examined the variables’ degradation rate. When upstream and downstream variables fluctuate with similar lifetimes, we find a greater probability of successfully inferring their edge direction. Furthermore, we have found a connection between the index of dispersion *D* and the ability to infer the direction of the interaction, where larger *D* yields a larger success ratio. Both degradation times and index of dispersion effect on the overall performance were previously demonstrated to be with similar conclusions as is given in [42]. Moreover, some of the network topological features such as the node degree were also being inspected. We have found that a lower incoming degree of the node provides better results in the inference of the direction. That is in agreement with previously published results that show that increasing the incoming degree requires more data points to gain the same quality of inference [27, 50].

To summarize, the method performs well where the joint distribution is well sampled. That is achieved when measuring sufficiently many data points such that they satisfactory capture the required statistical features. Nevertheless, as mentioned, here we provide only a basic algorithm that proves the principle that the direction of interaction is encoded within the stationary information. For future research we suggest examining further sophisticated approaches including stacking of methods and artificial neural networks [59–61].

## IV. METHODS

### A. Background Theory

The well-known Master equation for *V* variables yields at stationarity the global balance relation, means that for every variable *i*

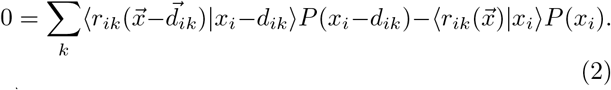

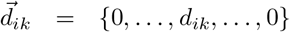 where the non-zero value is located in the *i*-th component. The above relation is derived from a simple summation of the stationary Master equation over all other variables, and has been derived and discussed previously [39– 42], see also derivation in SI. The angular brackets represent conditional means, 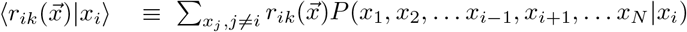 [62]. Eq. (2) can be wriiten as

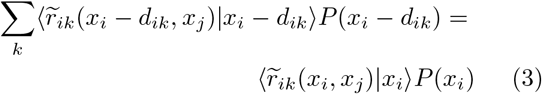

Where 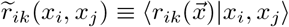. Our approach is based on *local* sensitivity of 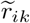 to *x*_*j*_, which aims to quantify the dependence of the dynamic of *x*_*i*_ on the level of *x*_*j*_. In particular we calculate *J*_*ij*_ *≡ ∂r*_*ik*_*/∂x*_*j*_ next to the probable abundance point. The quantities of |*J*_*ij*_| and |*J*_*ji*_| are then utilized as a classification feature. The nature of this comparison between |*J*_*ij*_| and |*J*_*ji*_|, and its role in classification would depend on the specific context or system. It is partially inspired by [56, 57] where both directions were quantified using regression and then assessed such that the more probable direction is chosen.

### B. Models

For numerical demonstrations we generate a random directed graph *Ĝ* which holds the topology of the network. Mathematically the graph is described by the adjacency matrix with elements *G*_*ij*_ = 1 where the state of *j* affects the dynamics of node *i*, i.e. *j → i*, and *G*_*ij*_ = 0 otherwise. We examined systems with Erdös Rényi random networks. In addition we examine more biologically realist system which is based on a subset of an *E. coli* gene regulatory network and was provided in the DREAM challenge [46–48].

For illustration, we use a typical reaction rate model where a molecule reaction can be either a reproduction 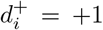 or degradation 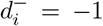. We specified that the production rate of *x*_*i*_, denoted as 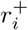, depends on other units states via the reaction network and that each molecule within the system degrades independently with a typical degradation rate. Hence, Eq. (2) can be written as

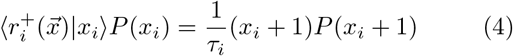

where 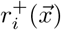 encodes all incoming edges (all units that affect the production rate of node *i*), and *τ*_*i*_ is the known lifetime of molecule *i*.

#### 1. Michaelis–Menten Regulatory Network

The birth rate of molecule *i* is

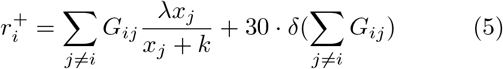

where for our simulation we choose *V* = 10 nodes, 10 directed edges, *λ* = 100 and *k* = 100. These commonly described production rates in biochemical reaction networks, where *λ* gives the maximal rate and *k* refers to the concentration which gives half of the maximal rate [63]. The degradation rates follow 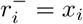 for all *i*.

#### 2. Coupled Goodwin Oscillators

Each Goodwin oscillator is given by three interacting variables (*x*_*i*_, *y*_*i*_, *z*_*i*_) which are coupled through the *y* variables. Their proliferation rates are given in the following:

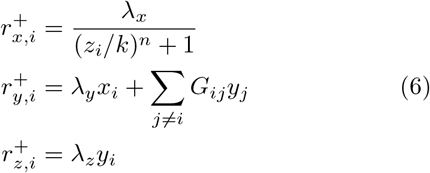

and death rates are 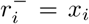 for all *i*. In here, the dynamic influences network within each oscillator is known, namely the relations within the triplet (*x*_*i*_, *y*_*i*_, *z*_*i*_) are given, we aim to infer only the coupling network between these triplets *G*_*ij*_. In the simulation presented in the main text, *Ĝ* has 8 edges and 8 nodes - where each node is a triplets, such as we have 24 fluctuated variables in total.

#### 3. Gene Regulatory Interactions in E. coli

As mentioned we also examine more biologically realist model which is based on a subset of an *E. coli* gene regulatory network and was provided in the DREAM challenge [46–48]. There, the variables *x*_*i*_ are described as continuous and possess small values. Here, for the discrete treatment of the model, we scaled the variables by 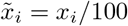, thus our *E. coli* gene regulatory network is thus defined and simulated with the following rates:

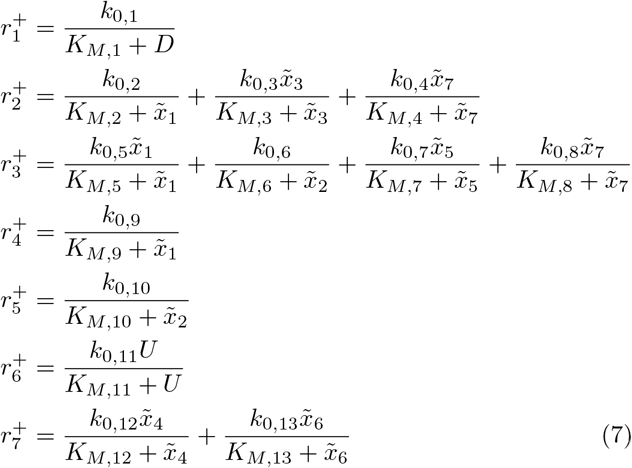

and for every *i*

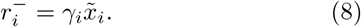

with parameters

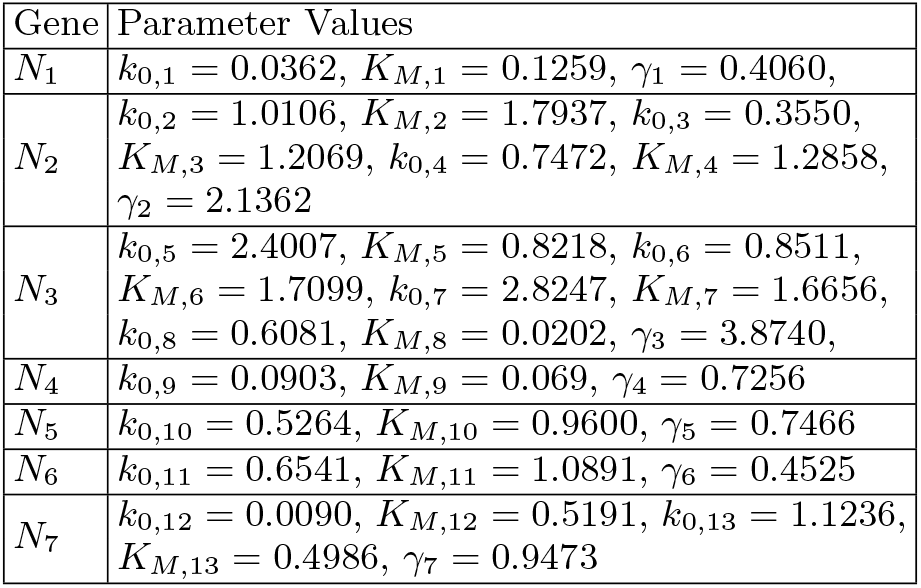

These parameters are taken from [46–48]. The ‘external disturbance’ is modeled by uniform distribution, means *D, U ∼* Uniform[0, 1].

