## supplemental text for "Determining Interaction Directionality in Complex Biochemical Networks from Stationary Measurements"

### SUPPLEMENTARY INFORMATION

#### Appendix A: Mathematical Background and Derivations

We consider a system with $V$ variables which pose the networks' nodes. Consider the multi-dimensional stationary master equation

$$0 = \sum_k \sum_{i=1}^{V} \left[ r_{ik}(\vec{x} - \vec{d}_{ik}) P(\vec{x} - \vec{d}_{ik}) - r_{ik}(\vec{x}) P(\vec{x}) \right]. \tag{A1}$$

$r_{ik}$ represents the $k$-th reaction rate of molecule $i$ and $\vec{d}_{ik} = \{0, \ldots, d_{ik}, \ldots, 0\}$ where the non-zero value is located in the $i$-th component. Without loss of generality, we index $x_i = x_1$. For every $i > 1$ we sum over all the possibilities of $x_i$, and find

$$0 = \sum_{x_2=0}^{\infty} \cdots \sum_{x_V=0}^{\infty} \left\{ \sum_k \sum_{i=1}^{V} \left[ r_{ik}(\vec{x} - \vec{d}_{ik}) P(\vec{x} - \vec{d}_{ik}) - r_{ik}(\vec{x}) P(\vec{x}) \right] \right\} \tag{A2}$$

Following the conditional probability relation $P(x_1, x_2 \ldots x_V)/P(x_1) = P(x_2 \ldots x_V | x_1)$ [1], we obtain $\sum_{x_2=0}^{\infty} \cdots \sum_{x_V=0}^{\infty} f(\vec{x}) P(\vec{x}) = \langle f(\vec{x}) | x_1 \rangle P(x_1)$ for an arbitrary function $f(\vec{x})$. Therefore, we can write

$$0 = \sum_k \left[ \langle r_{1k}(\vec{x} - \vec{d}_{1k}) | x_1 \rangle P(x_1 - d_{1k}) - \langle r_{ik}(\vec{x}) | x_1 \rangle P(x_1) \right] \tag{A3}$$

which recovers Eq. (1) in the main text. This relation though depends on the levels of all variables within the system via $P(x_1, x_2, \ldots x_V)$.

In the case presented in the main text, $d_{1k} = \pm 1$, then

$$\langle r_1^+(\vec{x}) | x_1 \rangle P(x_1) = \langle r_1^-(\vec{x} + \vec{e}_1) | x_1 + 1 \rangle P(x_1 + 1). \tag{A4}$$

We assume that the birth rate depends on the other molecules levels, and only self-degradation apply. Then

$$\langle r_1^+(\vec{x}) | x_1 \rangle P(x_1) = \sum_{x_2=0}^{\infty} \sum_{x_3=0}^{\infty} \cdots \sum_{x_V=0}^{\infty} r_i^+(x_2, x_3, \ldots x_n) P(x_2, x_3, \ldots x_V | x_1) P(x_1)$$

$$\langle r_1^-(\vec{x} + \vec{e}_1) | x_1 + 1 \rangle P(x_1 + 1) = \beta_1 (x_1 + 1) P(x_1 + 1). \tag{A5}$$

Thus, back to generality $x_i = x_1$, we obtain

$$\langle r_i^+(\vec{x}) | x_i \rangle P(x_i) = \beta_i (x_i + 1) P(x_i + 1). \tag{A6}$$

The latter can be written as

$$\langle r_i^+(\vec{x}) | x_i \rangle P(x_i) = \sum_{x_j=0}^{\infty} \langle r_i^+(\vec{x}) | x_j \rangle P(x_j | x_i) P(x_i) \equiv \sum_{x_j=0}^{\infty} \widetilde{r}_i^+(x_j) P(x_j | x_i) P(x_i)$$

$$= \langle \widetilde{r}_i^+(x_j) | x_i \rangle P(x_i) = \beta_i (x_i + 1) P(x_i + 1). \tag{A7}$$

From the above relation one can determine $\widetilde{r_i}^+(x_j)$ as a function of $x_j$ from stationary distribution $P(x_i, x_j)$ solely as shown in [2].

#### Appendix B: A Comment About Non-Stationary Processes

Consider the general time-dependent Master equation

$$\partial_t P_n(t) = \sum_{k \neq n} a_{kn} P_k(t) - \sum_{k \neq n} a_{nk} P_n(t). \tag{B1}$$

Applying time averaging on both sides yields

$$\frac{P_n(T) - P_n(0)}{T} = \sum_{k \neq n} a_{kn} \frac{1}{T} \int_0^T P_k(t) - \sum_{k \neq n} a_{nk} \frac{1}{T} \int_0^T P_n(t) \tag{B2}$$

$$\frac{P_n(T) - P_n(0)}{T} = \sum_{k \neq n} a_{kn} \overline{P_k}(T) - \sum_{k \neq n} a_{nk} \overline{P_n}(T), \tag{B3}$$

which presents a similar form to stationary Master equation Eq. (A1) in cases where $\lim_{T\to\infty} \frac{P_n(T)-P_n(0)}{T} = 0$ and the probability distributions are replaced with their time averaged ones.

#### Appendix C: Simulations Details

After presenting our mathematical framework, we examine how useful is our method to analyze noisy data from finite measurements. The time propagation simulations, that had been done using the Gillespie algorithm, are for the purpose of mimicking various experimental scenarios.

From the stochastic processes described in Methods, we measure the stationary probability density function (PDF) $P(x_i, x_j)$ as explained in the following. We first construct a long time realization of the variables under consideration. Such a realization is routinely constructed using the Gillespie algorithm [3]. In our simulation, we use $2 \cdot 10^7 \cdot V$ time steps, where $V$ is the number of variables simulated. We assume that the distribution obtained from that very-long realization is thus exact. Then, we sample from the "exact" PDF a given number of data points $N$ corresponding to their probability.

Here we use the same algorithm used in [2] to infer $\widetilde{r}_i^+(x_j)$ from $P(x_i, x_j)$. There, the authors found the birth rate $\widetilde{r}_i^+(x_j) \equiv \vec{f}$ by finding

$$\underset{f}{\operatorname{argmin}} \left\{ \left| \hat{G}\vec{f} - \vec{h} \right|^2 + \epsilon \left| \hat{\Gamma}\vec{f} \right|^2 \right\} \qquad \text{subject to } \vec{f} \geq 0 \tag{C1}$$

where we defined $G_{mn} \equiv P(x_i = m, x_j = n)$, $f_n \equiv \widetilde{r}_i{}^+(x_j = n)$ and $h_m \equiv \beta_i(m+1)P(x_i = m)$. The additional regularization matrix $\Gamma_{m,n} \equiv \delta_{m,n} - \delta_{m,n+1} + \delta_{m,n+2}$ penalizes non-smooth functions with regularization constant $\epsilon = N^{-1/2}$ where $N$ is the number of sampled data points, see [2].

#### Appendix D: The Distribution of the Conditional Expectation

Here we use examples where the proliferation rate may depends on other molecules level $r_i^+(\vec{x})$, and death rate are self govern degradation $r_i^-(x_i) = \beta_i x_i$ As mentioned shown

$$\langle r_i^+(\vec{x}) | x_i \rangle = \frac{\beta_i(x_i+1)P(x_i+1)}{P(x_i)}. \tag{D1}$$

We note that the *measured* $P(x_i)$, marked as $\hat{P}(x_i)$, is essentially a random number where noise is induced by the finite sampling. The probability that from $N$ samples, we find $k$ realizations with level $x_i$ of molecules from type $i$ is given by

$$\mathrm{PDF}(k) = \binom{N}{k} P(x_i)^k (1 - P(x_i))^{N-k}. \tag{D2}$$

Then, by a simple transformation, we obtain that the estimated (measured) value $\hat{P}(x_i)$ is distributed via

$$\mathrm{PDF}\left[\hat{P}(x_i) = \frac{k}{N}\right] = \frac{P(x_i)^{N\hat{P}(x_i)} \left[1 - P(x_i)\right]^{N - N\hat{P}(x_i)}}{B\left[N\hat{P}(x_i) + 1, N - N\hat{P}(x_i) + 1\right]}, \tag{D3}$$

where $B[\alpha, \beta]$ refers to the Beta function. When $N \gg 1$, $NP(x_i) \gg 1$ and $N[1 - P(x_i)] \gg 1$ we find

$$\hat{P}(x_i) \sim \mathcal{N}\left[P(x_i), \frac{P(x_i)\,(1 - P(x_i))}{N}\right] \tag{D4}$$

which is the Gaussian distribution with mean $\langle \hat{P}(x_i) \rangle = P(x_i)$ and variance $\mathrm{Var}\left[P(\hat{x_i})\right] = P(x_i)\left[1 - P(x_i)\right]/N$.

Now, the distribution of the evaluated $z = \langle r_i^+(\vec{x}) | x_i \rangle$ is controlled by the ratio of two random numbers. In the case where both the numerator and the denominator are Gaussian distributed, we can use the transformation

$$t \approx \frac{\mu_1 z - \mu_2}{\sqrt{\sigma_1^2 z^2 - 2\rho\sigma_2\sigma_1 z + \sigma_2^2}}. \tag{D5}$$

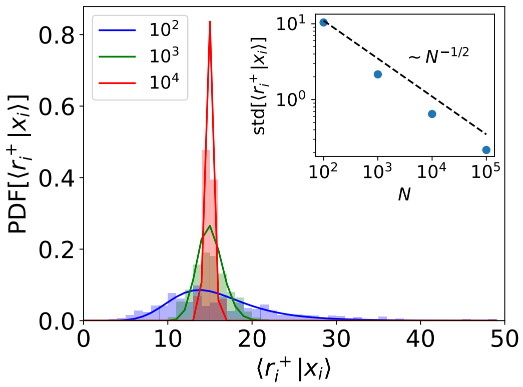


FIG. 1. The distribution PDF $\left[z = \langle r_i^+(\vec{x}) | x_i \rangle\right]$ evaluated at $x_i = \mathrm{ArgMax}[P(x_i)]$. It is approximated as correlated Gaussian ratio distribution, and evaluated using Geary-Hinkley transformation Eq. (D5). Inset: The measured standard deviation for large $N$ falls as $N^{-1/2}$.

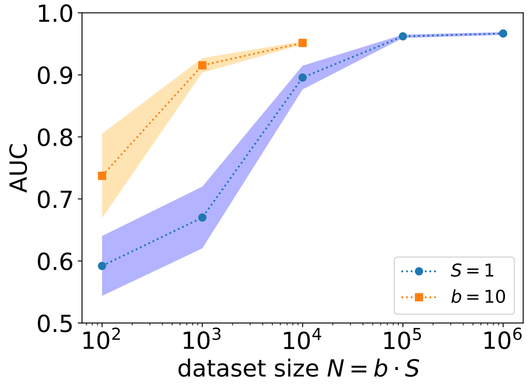


FIG. 2. Using small batches might be beneficial, especially for small $N$. The results are given from the Goodwin oscillators model.

with $\mu_1 \equiv \langle \hat{P}(x_1) \rangle$, $\mu_2 = \langle \beta_i(x_i+1)\hat{P}(x_i+1) \rangle$,$\sigma_1^2 \equiv \mathrm{Var}\left[\hat{P}(x_i)\right]$, $\sigma_2^2 \equiv \mathrm{Var}\left[\beta_i(x_i+1)\hat{P}(x_i+1)\right]$ and $\rho$ is the Pearson correlation coefficient between the numerator and the denominator. Then $t \sim \mathcal{N}(0,1)$ following the Geary-Hinkley theorem [4, 5]. Note that PDF $\left[z = \langle r_i^+(\vec{x}) | x_i \rangle\right]$ is a wide distribution, yet simulation results suggest that when the distributions of the numerator and denominator are sufficiently narrow, the standard deviation of $z$ decreases as $1/\sqrt{N}$, where $N$ is the number of samples. This is true only where $\hat{P}(x)$ is Gaussian distributed, i.e. where $x$ is in the central part of $P(x)$. In Fig. 1 we illustrate the statements given above.

#### Appendix E: Dependence of the number of subdivisions and batch size of the data

Consider that the total data length is $N$, which may represent the total number of samples or the data points. Then we divide this set into $S$ equal-sized sets, namely batches, where the size of each batch is $b$ (note that $N = S \cdot b$). With the same amount of data points $N$, one may find different quality of estimation depending on the number of the subdivisions $S$ and the batch size $b$. We have found by simulation that for a given $N$, there is an advantage of taking small batches with $b = 10$ samples, see Fig. 2 . This phenomenon might be partially explained by the following. Too low batch size $b$ might be associated with underfitting - where the amount of information within each batch is insufficient to make any inference. Yet, there is an essential non-linear relation between the sampling error of $P(x_i, x_j)$ and the quality of estimation (see previous section).

#### Appendix F: Index of Dispersion

In cases where $x_i$ is independent on other variables; means independent proliferation $r_i^+ = \text{const.} = c_i$ with degradation $r_i^- = \beta_i x_i$, we find $P(x_i) = e^{-c_i/b_i}(c_i/b_i)^n/x_i!$. The latter is a Poisson distribution with index of dispersion $\text{var}(x_i)/\langle x_i \rangle \equiv D = 1$.

The stationary Master equation where $r_i^+(\vec{x})$ may generally depends on other molecules' levels and degradation $r_i^-(x_i) = \beta_i x_i$ is

$$0 = \sum_{i=1}^{V} \left[ r_i^+(\vec{x} - \vec{e}_i) P(\vec{x} - \vec{e}_i) - r_i^+(\vec{x}) P(\vec{x}) + \beta_i (x_i + 1) P(\vec{x} + \vec{e}_i) - \beta_i x_i P(\vec{x}) \right] \tag{F1}$$

where we simply used Eq. (A1) with two possible reactions for each molecules $d_{ik} = d_i^{\pm} = \pm 1$. Multiply by $x_1$ and sum over all possible states $\sum_{x_1=0}^{\infty} \cdots \sum_{x_V=0}^{\infty}$ yields

$$0 = \sum_{x_1=0}^{\infty} \cdots \sum_{x_V=0}^{\infty} \sum_{i=1}^{V} x_1 \left[ \underbrace{r_i^+(\vec{x} - \vec{e}_i) P(\vec{x} - \vec{e}_i) - r_i^+(\vec{x}) P(\vec{x})}_{\text{I}} + \underbrace{\beta_i (x_i + 1) P(\vec{x} + \vec{e}_i) - \beta_i x_i P(\vec{x})}_{\text{II}} \right]. \tag{F2}$$

Then

$$\sum_{x_1=0}^{\infty} \cdots \sum_{x_V=0}^{\infty} \sum_{i=1}^{V} x_1 \cdot (\text{II}) = \sum_{x_1=0}^{\infty} \cdots \sum_{x_V=0}^{\infty} \sum_{i=1}^{V} x_1 \beta_i (x_i + 1) P(\vec{x} + \vec{e}_i) - x_1 \beta_i x_i P(\vec{x}) = \beta_1 \langle x_1 \rangle \tag{F3}$$

$$\sum_{x_1=0}^{\infty} \cdots \sum_{x_V=0}^{\infty} \sum_{i=1}^{V} x_1 \cdot (\text{III}) = \langle r_1^+(\vec{x}) \rangle$$

Thus we find that $\beta_1 \langle x_1 \rangle = \langle r_1^+(\vec{x}) \rangle$. Similarly, we can multiply by $x_1^2$ and find

$$\sum_{x_1=0}^{\infty} \cdots \sum_{x_V=0}^{\infty} \sum_{i=1}^{V} x_1^2 \cdot (\text{II}) = \beta_1 \langle x_1 \rangle - 2\beta_1 \langle x_1^2 \rangle \tag{F4}$$

$$\sum_{x_1=0}^{\infty} \cdots \sum_{x_V=0}^{\infty} \sum_{i=1}^{V} x_1^2 \cdot (\text{III}) = 2 \langle x_1 r_1^+(\vec{x}) \rangle + \langle r_1^+(\vec{x}) \rangle$$

Using the above we find

$$\begin{aligned} 2\beta_1 \langle x_1^2 \rangle &= \beta_1 \langle x_1 \rangle + 2\langle x_1 r_1^+(\vec{x}) \rangle + \langle r_1^+(\vec{x}) \rangle \\ 2\beta_1 \langle x_1^2 \rangle - 2\beta_1 \langle x_1 \rangle^2 &= \beta_1 \langle x_1 \rangle + 2\langle x_1 r_1^+(\vec{x}) \rangle + \langle r_1^+(\vec{x}) \rangle - 2\langle r_i^+(\vec{x}) \rangle \langle x_1 \rangle \\ \beta_1 \text{var}(x_1) &= \beta_1 \langle x_1 \rangle + \text{cov}\left(x_1, r_i^+(\vec{x})\right) \end{aligned} \tag{F5}$$

Hence we obtain

$$D \equiv \frac{\text{var}(x_1)}{\langle x_1 \rangle} = 1 + \frac{\text{cov}\left(x_1, r_1^+(\vec{x})\right)}{\langle r_1^+(\vec{x}) \rangle}. \tag{F6}$$

Where the index of dispersion $D$, also called the Fano factor, effectively quantifies the contribution of the correlation between $r_i^+(x_j)$ and $x_i$, to the variability of $x_i$ as was previously shown in [6]. In cases where $x_i$ is weakly depend on other variables; means $r_i^+ \approx \text{const.}$ we we find $D \approx 1$.

#### Appendix G: Network Topological Properties

We survey several topological properties of the network and their effect on the overall performance. In particular, we examine the local topological properties for the edge $x_i \to x_j$, which includes the in- and out-degree of both $x_i$ and $x_j$, see Fig. Our simulations do not significantly indicate the influence of local degrees on the performance, yet a slight dependence observed in the in-degree of $x_i$ as shown in the main text.

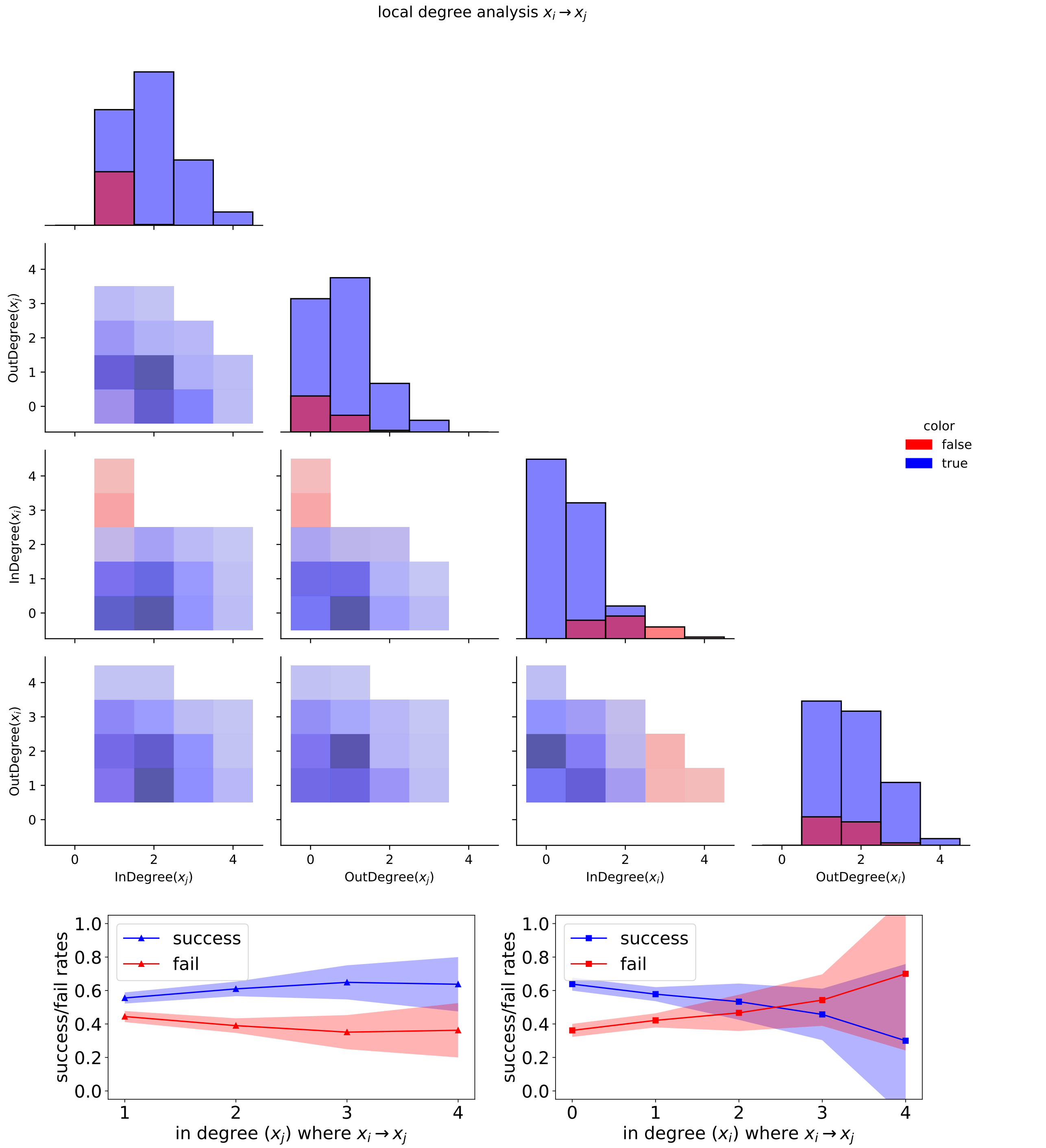


FIG. 3. local degree analysis on the overall performance. Upper panel presents results between pairs of features (=degrees). In the lower row we show the results for the in-degree of $x_i$ and $x_j$ (the former is the same as in the main text).

#### Appendix H: Classification Features

##### 1. Considered Attributes

We have tested three attributes, i.e. quantities, which aim to capture the strength of association from one variable to the other. The first one is based on the sensitivity analyses of the dynamic rate. There the attribute is the slope

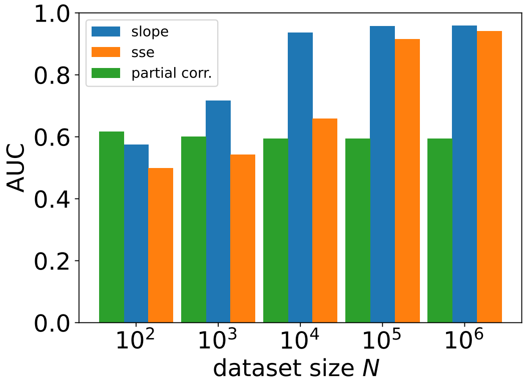


FIG. 4. Examination of classification features. We found that the classifying quantity which is the slope at some $\vec{x}^*$, namely $|J_{ij}|$, out-performs the SSE-based feature especially in low $N$. In addition, we have recorded all other variables within the network to evaluate the partial correlation.

$|J_{ij}|$ - which quantifies the changes in the rate of $x_i$, i.e. $\tilde{r}_i(x_j)$, while changing the level of another variable $x_j$. That attribute was used throughout the main text and the SI up to here.

Another classification attribute that was examined is described in the following. In [7, 8] the authors suggested using the Minimum Description Length (MDL) principle which is described as a proxy for the Kolmogorov complexity. There they define a quantity $\Delta_{x\to y} = [L(x) + L(y|x)] / [L(x) + L(y)]$ and $\Delta_{y\to x}$ is defined analogously. Then infer $x \to y$ , if $\Delta_{x\to y} < \Delta_{y\to x}$ holds up to an additive constant. The MDL is approximately the sum-of-squared errors (SSE). Here we quantify the SSE where assuming that $\tilde{r}_i(x_j)$ indeed depends on $x_j$, the SSE where assuming $\tilde{r}_i$ is independent of $x_j$, as well as the SSE of $\tilde{r}_j(x_i)$ and $\tilde{r}_j$. The so-called SSE attribute presented in Fig. 4

The last classification attribute we examine is the partial correlation, which disadvantageously requires recording all variables within the system. We have found that the partial correlation feature exceeds the completely random classifier, i.e. it exceeds AUC=0.5. However, it does not necessarily yield better results and thus is not preferred over the other classification features, particularly for relatively high $N$. The results are presented in Fig. 4 with an agreement with [9].

##### 2. Feature Engineering - Oriented Graphs

Hitherto, we consider the undirected edge $(i, j)$ is given with its corresponding stationary joint distribution $P(x_i, x_j)$ and aim to determine its binary direction, which means either $i \to j$ or $j \to i$. In graph theory that task is called 'graph orientation', which aims to assign a single direction to every edge in an undirected graph.

Let consider the attribute $J_{ij} \equiv \frac{d\tilde{r}_i(x_j)}{dx_j}|_{x_j=x_j^*}$. Thus, for a given edge we infer its direction as follows; if $|J_{ij}| > |J_{ji}|$ an arrow from $j$ to $i$ is drawn, and vise-versa - if $|J_{ij}| \geq |J_{ji}|$ we infer an arrow from $i$ to $j$. It gives that the classification feature is in effect the quantity $|J_{ij}| - |J_{ji}|$. Throughout the main text and the SI up to here, the results are presented for the binary classification, with a preprocess of the features engineering as described above.


[1] F. Edition, A. Papoulis, and S. U. Pillai, *Probability, random variables, and stochastic processes* (McGraw-Hill Europe: New York, NY, USA, 2002).

[7] A. Marx and J. Vreeken, Telling cause from effect using mdl-based local and global regression, in *2017 IEEE inter-*

*national conference on data mining (ICDM)* (IEEE, 2017) pp. 307–316.

[8] P. Blöbaum, D. Janzing, T. Washio, S. Shimizu, and B. Schölkopf, Analysis of cause-effect inference by comparing regression errors, PeerJ Computer Science **5**, e169 (2019).

[9] M. Nitzan, J. Casadiego, and M. Timme, Revealing physical interaction networks from statistics of collective dynamics, Science advances **3**, e1600396 (2017).
